# Improved localizers and anatomical images to enable phosphorus magnetic resonance spectroscopy of liver metastasis at 7T

**DOI:** 10.1101/315572

**Authors:** Debra Rivera, Irene Kalleveen, Catalina Arteaga de Castro, Hanneke van Laarhoven, Dennis Klomp, Wybe van der Kemp, Jaap Stoker, Aart Nederveen

## Abstract

Phosphorus spectroscopy (31P) at 7T (300 MHz) enables clinically-relevant spatial resolutions and time scales with high potential for monitoring response to cancer treatment. However, at 7T collecting a radiological-grade anatomical image of the liver—which is required for performing localized 31P spectroscopy—presents a challenge. Unlike lower field-strength scanners, there is no body coil in the bore of the 7T and despite inadequate penetration depth (<10 cm), surface coils are the current state-of-the-art for acquiring anatomical (1H) images. Therefore, thus far, high field 31P spectroscopy has been limited to diffuse liver disease. However, the use of antennas enable improved penetration depths at 300 MHz, and when combined with parallel transmit, can enable body imaging at 7T. We have developed a protocol for imaging liver metastases of patients using parallel transmit and 31P spectroscopy at 7T. We used a custom-made liver coil consisting of eight 30-cm dipole antennas tuned to the proton (300 MHz) frequency, and two partially overlapping 20-cm-diameter loops tuned for 31P (120 MHz). The field of view afforded by the two antennas underneath the 31P loops is not sufficient to image the complete boundaries of the liver for chemical shift imaging (CSI) planning and region-of-interest-based B0 shimming. The liver and full axial slice of the abdomen was imaged with eight transmit/receive antennas using parallel transmit B1-shimming to overcome image voids. Through the use of antennas we overcome the challenges for multi-parametric body imaging, and can begin to explore the possibility of monitoring the response of patients with liver metastasis to cancer treatments.

**ABBREVIATIONS:**

(PDE): Phosphodiester
(GPE): Glycerophosphoethanolamine
(GPC): Glycerophosphocholine
(PME): Phosphomonoesther
(PC): Phosphocholine
(PE): Phosphoethanalomine
(PI): Inorganic Phosphate
(PCR): Phosphocreatine
(PTC): Phophotidylcholine
(CSI): Chemical Shift Imaging
(GE): Gradient Echo
(L): Left
(R): Right
(H): Head
(F): Foot
(A): Anterior
(P): Posterior
(TR): Repetition Time
(TE): Echo Time

## Introduction

Thus far, high field phosphorus (31P) spectroscopy in humans has been limited to diffuse liver disease, due to the difficulty of obtaining proton images to prepare the phosphorus chemical shift imaging (CSI). Ultra-high field (> 4T) magnetic resonance imaging brings phosphorus spectroscopy (31P) to clinically relevant spatial and temporal resolutions and allows for differentiation of relevant metabolites, opening up the possibility of 31P spectroscopy as a non-ionizing method of metabolic tracking of tumor response to therapeutics in patients.^1^ The liver is a frequent site for metastases of solid tumors, such as breast, colon, oesophagogastric, and pancreatic cancers.^2,3^ Once metastasized, most cancers are incurable, although overall survival and quality of life may be extended by systemic treatment.^4^ Unfortunately, response to treatment varies greatly between patients, even when suffering from the same primary tumor.^5^ New tools to measure or predict treatment response shortly after start of treatment are urgently needed to prevent long-term exposure of patients to ineffective treatments. ^6^ Phosphorus spectroscopy provides a multitude of information on cancer metabolism, and the local environment such as: phospholipid metabolism (phosphoethanolamine (PE), phosphocholine (PC), glycerophosphoethanolamine (GPE) and glycerophosphocholine (GPC)), cellular energetics, and intracellular and extracellular pH (via inorganic phosphate (Pi)).^7^ These may all be related to (early) treatment response.^7,8^ In the preclinical setting, 31P spectroscopy is already a key tool for elucidating cancer metabolism and quantitative evaluation of new treatments.^9^ As such, clinical 31P spectroscopy is of great interest as a potential tool for tracking treatment response.^6^

At 7T (300 MHz) collecting a proton image of the liver presents a challenge as the wavelength for proton imaging at 300 MHz approaches the dimensions of the abdomen.^10,11^ Therefore, conventional approaches using volume coils (e.g. birdcage) or loop coils for acquiring scout images for preparing 31P scans or anatomical images on which to overlay spectra results (localizer images) no longer give sufficient penetration depth for proton imaging at high field.^12,13^ There is no body coil in the bore of the 7T, nor are receive loops present in the bed.

Antennas propagate electro-magnetic fields directly into the tissue, and thus enable improved penetration depths over loops and microstrip elements, overcoming many of the challenges associated with body imaging at 300 MHz.^14,15^ In addition to poor penetration depth, at high field destructive interference causes inhomogeneity of the transmit field, ^16^ which can be minimized by a technique known as B1+ shimming. In B1+ shimming, the transmit profiles of each element are mapped in order to determine the required input phase (and optionally magnitude) of the excitation pulses sent to individual transmit elements in order to reduce destructive interference in the region of interest.^10,16,17^ In practice, B1+ shimming requires a system capable of independent control of multiple transmit channels that can operate simultaneously, namely a parallel-transmit system.

Motivated by the potential of 31P ultra-high-field spectroscopy as a means of optimizing treatments on an individual patient level, and recent successes with antenna elements for body imaging at 300 MHz, we set out to develop a hardware solution and imaging protocol to obtain phosphorus spectroscopy and corresponding parallel-transmit anatomical images for cancer patients with metastatic disease in the liver. Here we present the use of antennas to improve penetration depth for multi-nuclear spectroscopy, as an improvement over the common practice of using loops for obtaining localizers for phosphorous spectroscopy,^18,19^ and demonstrate the need for the combination of parallel transmit with phosphorus spectroscopy (Fig. 1). We provide aggregate data (n=9) demonstrating the limiting coverage of the two antennas underneath the 31P loops, and present 3D CSI 31P spectra of a patient with metastatic liver cancer superimposed on an anatomical image collected with a parallel-transmit system that combines the two elements with another six elements.

**Fig. 1.**
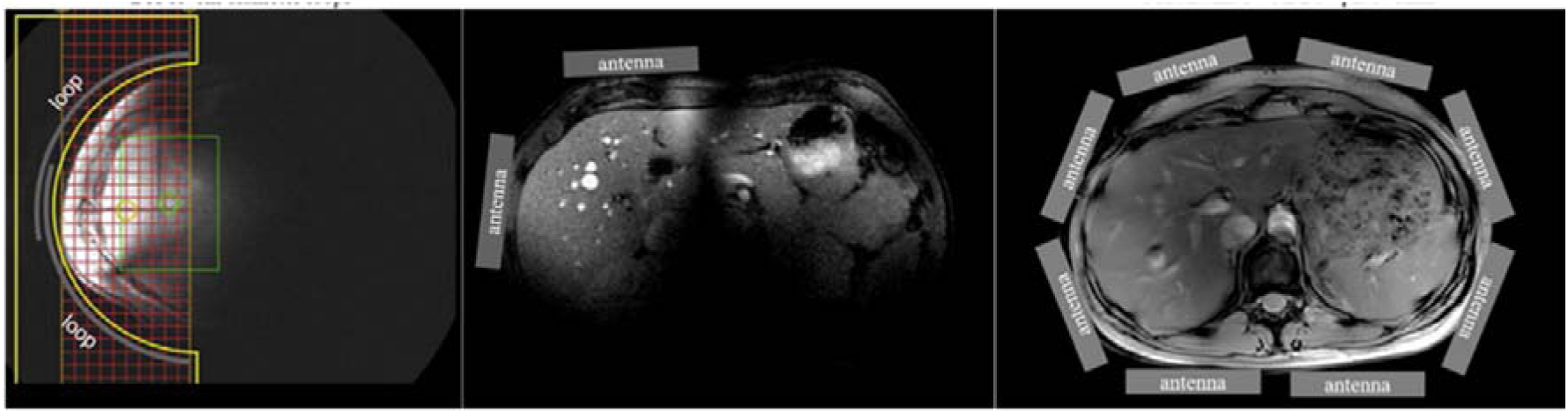
Two overlapping loops are the state-of-the-art for localizer images for multinuclear experiments at 7T (left), adapted from Runge et al. By using two antennas, the coverage of the liver is extended to include the anatomical left and right hemi-livers (center). Eight antennas used in combination with parallel-transmit (B1+) phase-shimming provides full coverage of the liver and the entire cross-section of the abdomen (right).

## Methods

A custom liver coil (MR Coils, BV, Zaltbommel, The Netherlands) was built for use with a 7T Philips system (Philips, Cleveland, Ohio, USA) consisting of eight fractionated dipole antennas (30 cm from end to end of the conductors—according to Raaijmakers et al. for transmit and receive of the proton (300 MHz) signal,^20^ and two partially overlapping loops (20 cm in diameter) for transmitting and receiving the 31P (120 MHz) signal. The study was approved by the internal review board, and written informed consent was obtained from all participants. A phantom filled with a 55% solution of alcohol in water with 4.8 g/L salt (permittivity of 36 and a conductivity of 0.43 S/m) was used to demonstrate penetration depth of a single transmit element.

The multi-nuclear experiment and associated proton imaging were conducted with two (out of eight) antennas (300 MHz) and the two overlapping loops (120 MHz). For each frequency, the pairs of elements were combined with quadrature hybrids (+90-degree for transmit, -90-degree for receive). Power was optimized for the 31P sequence by using a flip-angle sweep to identify the maximum signal intensity in a phantom (detailed above) and verified in a participant. Similarly, the optimal power level was found for the proton antennas as evaluated in a phantom and verified in vivo, to verify that the desired flip-angle was in agreement with the actual flip-angle. Adiabatic pulse-acquire scans were performed for the multinuclear experiments; therefore, we did not further optimize the transmit power.

As of yet, no commercially available 7T body MR system can simultaneously support parallel transmit proton imaging and multi-nuclear experiments. Therefore, the scanner was rebooted mid-session in order to access the transmit chain for the multi-nuclear experiment (Classic mode) for half of the examination, and the parallel-transmit capable transmit chain (Multix mode) during the other half of the experiment.

### Classic-mode protocol

Proton (^1^H) localizer images were obtained using two antennas in quadrature for ROI-based B0-shimming (2^nd^-order) with a gradient echo MRI (parameters listed below). Phosphorus spectroscopy data was acquired with quadrature-combined overlapping loops. A pulse-acquire sequence (block pulse, 30° flip angle, 2048 samples, 8192 Hz BW, TR 1 s, 24 averages) was used to set the center frequency to correspond with PE. We obtained 3D CSI scans with field of views (FOV) of 320 mm × 320 mm × 320 mm TR/TE 700/0.4 ms and 40 mm isotropic voxels, with a total imaging time of 5 minutes per average. An adiabatic half passage RF pulse was used for pulse-acquire for the 3D CSI with a tanh/tan shape, 2-ms-pulse duration, and maximum frequency modulation of 21 kHz (174 ppm).

### Multix-mode protocol

The parallel transmit protocol was adapted from previous work using the same antenna elements for imaging the prostate, pancreas, and kidneys (e.g.,^21^). B1 and B0 shimming were performed using an image-based shim tool (MR Code, B.V., Zaltbommel, The Netherlands). Region-of-interest B0 shimming (2^nd^ order) was performed on a reference scan, and B1-shimming was conducted by mapping the B1 of each transmit antenna and calculating the relative phase offsets using a third-party software application running on the scanner host (MR Code, B.V., Zaltbommel, The Netherlands).

Images were acquired with a gradient echo using TR/TE 4.97/2.42 ms, 40° flip angle, 300 mm × 330 mm × 330 mm field of view, and voxels of 0.8 mm × 0.9 mm × 5 mm. Dixon images were collected in both modes (multi-slice gradient echo, 20 slices, TE1/TE2/TR 2.65/3.15/10 ms, 15° flip angle, 250 mm × 300 mm × 80 mm field of view, 276 × 276 encoding in 4-mm-thick slices, 3 averages, 109s acquisition).^22^ The field of view for obtaining water images with the Dixon method in the patient was adjusted to 234 mm × 359 mm × 80 mm.

Spectra voxels were selected from 3DiCSI (Hatch Center for MR Research Columbia University) and Hamming filtered, uploaded into JMRUI^23^and filtered with a 40 Hz Lorentzian, zero-filled to double the number of points (from 512 to 1024), zero-order phase corrected, and 1^st^-order (0.4 ms) phase corrected.

While considerable work has been done to characterize the use of eight-antenna arrays for body imaging, this is the first time that two antennas have been used as a means of localizing an anatomical target for multinuclear spectroscopy. Therefore, we characterized the depth of the sensitivity profile (calculated as 1.5 times greater than mean background signal) for nine participants along three axis: anterior/posterior (A/P), left/right (L/R), and head/foot (H/F). In order to evaluate the needed depth to provide full coverage of the liver, we characterized the depth of the liver along the same three axis for young (<40 years) healthy volunteers (2 male, 3 female). All length measurements are reported as the mean accompanied by standard deviation of the sample population in parentheses.

## Results

We have successfully combined multinuclear spectroscopy 8-channel antennas and parallel-transmit anatomical data in a single exam. The liver and full axial slice of the abdomen can be visualized with eight transmit/receive antennas (Fig. 2). Two antennas with fixed-transmit phases allowed the Couinaud system left and right hemi-livers to be visualized (Fig. 3). However, the liver-coverage achieved by two antennas was insufficient to visualize the boundaries of four out of nine (44%) of the healthy volunteers scanned. Aggregate data compare penetration depth (n=9) with liver depth (n=5 as four of the livers were not fully visualized) for each direction (Table 1). The penetration depth in the R/L and H/F direction exceed the needed depths for visualizing the liver, although the A/P penetration is approximately equal to the liver depth, and is therefore a limitation. The coverage in the H/F axis exceeds the needed coverage by two-fold. A gradient echo image demonstrates that a single antenna can provide signal along the entire depth of a 20-cm deep phantom (Fig. 4).

**Table 1.** Measurements of penetration depths and needed coverage for liver in the anterior/posterior (A/P), right/left (R/L), and head/foot (H/F) directions. Aggregate values are displayed in the table as mean (std) across participants. *Four of the nine participants were excluded in estimating the needed coverage, as the A/P boundary was insufficiently imaged.

|  | A/P [cm] | R/L [cm] | H/F [cm] |
| --- | --- | --- | --- |
| penetration (n=9) | 16.4 (1.5) | 27.9 (2.7) | 30.0 (2.7) |
| needed coverage (n=5*) | 14.8 (1.4) | 20.0 (2.4) | 15.1 (1.3) |

**Fig. 2.**
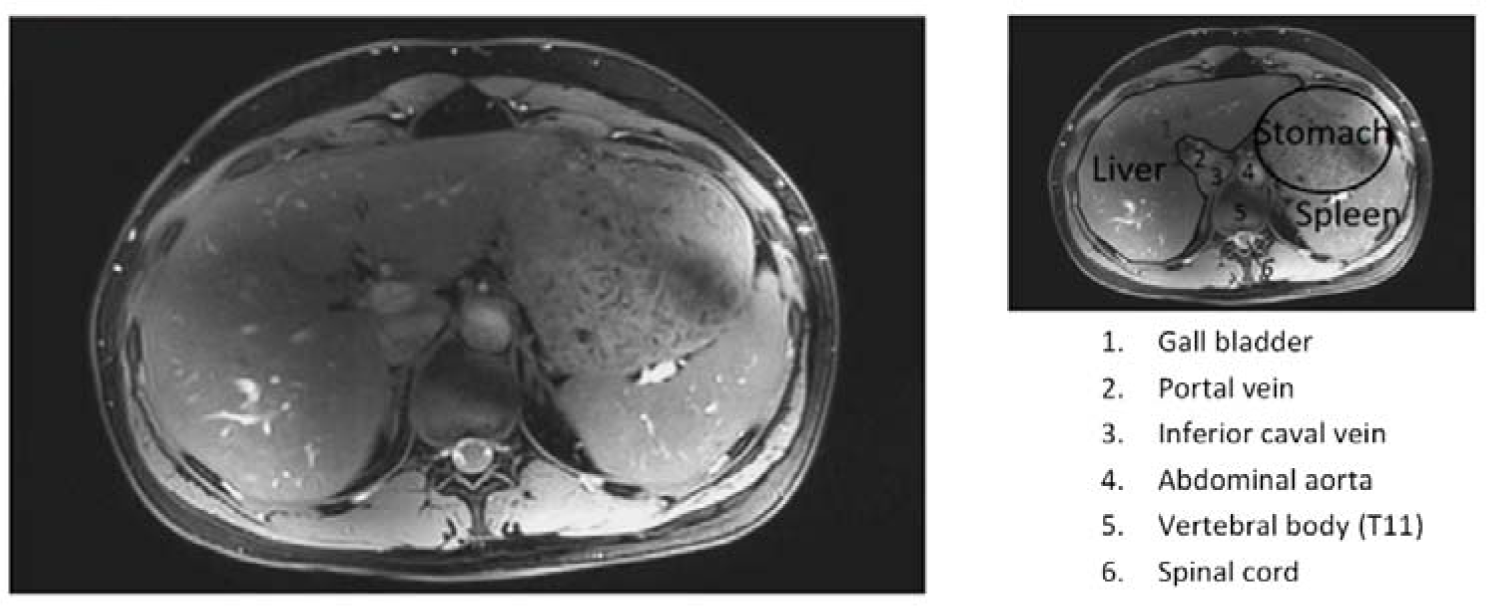
Dixon image of a healthy female participant (body mass index = 23, age = 38) using eight antennas and parallel transmit. Anatomical labels are provided on the identical image as to not obscure features.

**Fig. 3.**
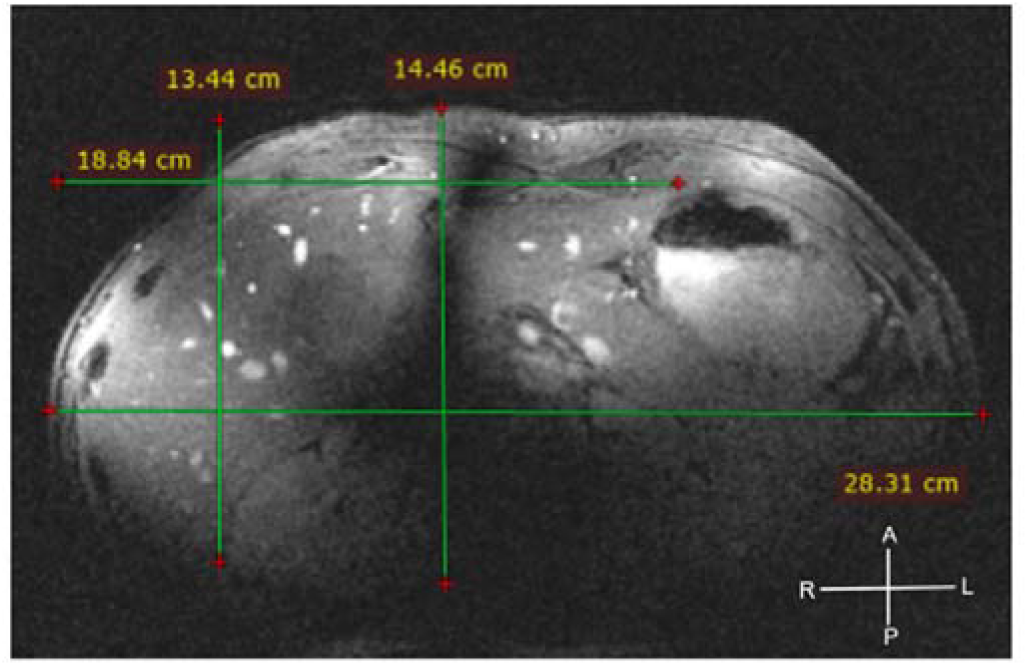
Measurements of penetration depths and needed coverage for liver imaging superimposed on a gradient echo image from a participant.

**Fig. 4.**
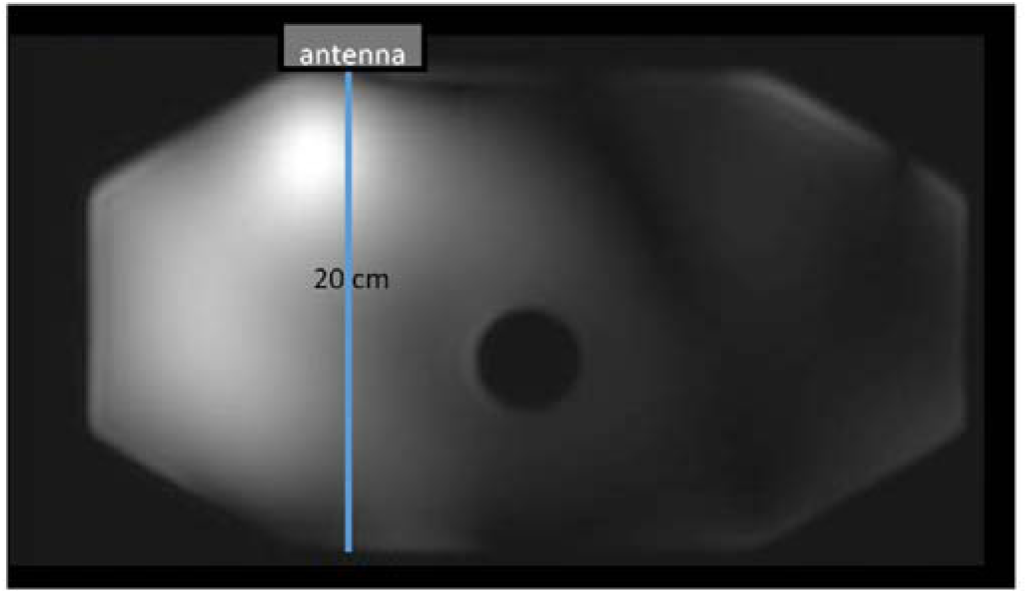
A gradient echo image created with a single antenna element perpendicular to the blue line, showing signal penetration along the full thickness of a 20-cm phantom (A/P direction).

While the right and left hemi-livers (Couinaud system) can be imaged with two antennas combined in quadrature, full visualization of the liver is not possible due to voids in the image (observable in Fig. 3). Parallel transmit with eight antennas allows for B1-shimming to overcome these image voids. Fig. 5 presents the identical slice performed with and without B1-phase shimming. The voids due to destructive interference are outlined in blue to aid comparison between the images. The eight antennas provide coverage of the complete region of interest (Fig. 6).

**Fig. 5.**
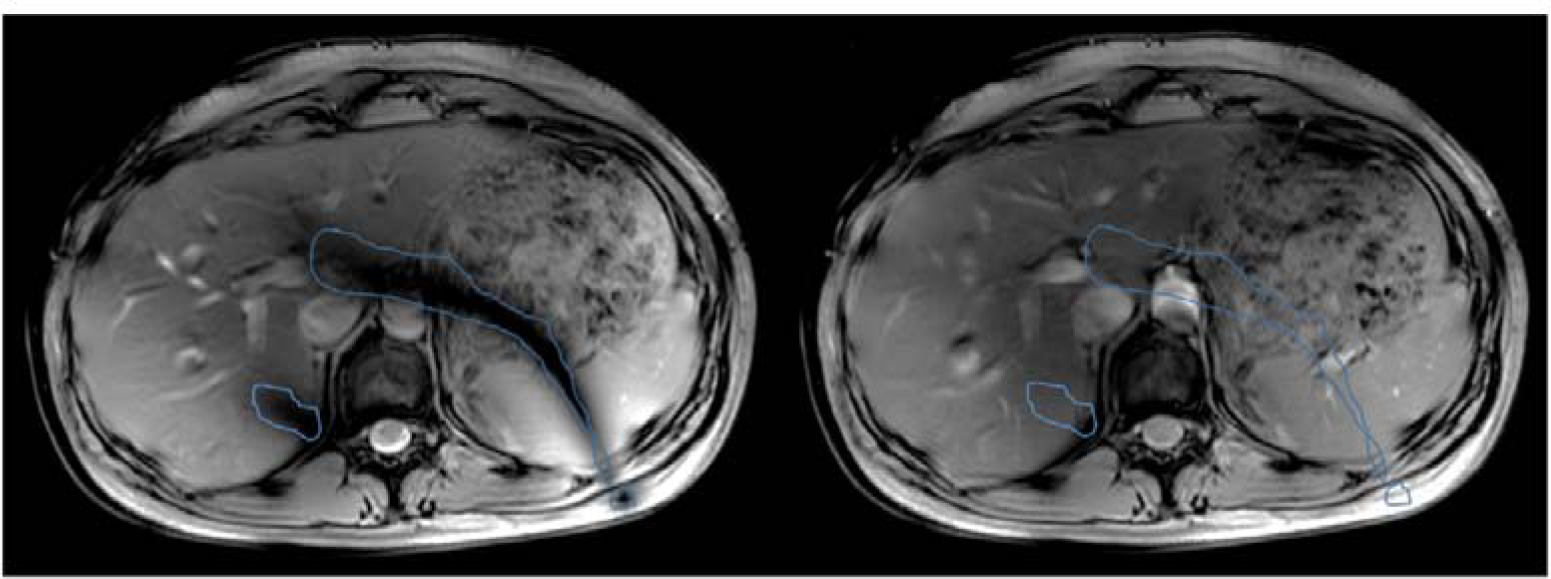
Radiological-grade anatomical imaging is possible with parallel-transmit B1-shimming using eight antennas. The signal voids (outlined in blue) due to destructive interference are present before B1-shimming (left) and gone after (right) adjusting relative phases of the transmit pulses (B1+ phase shimming).

**Fig. 6.**
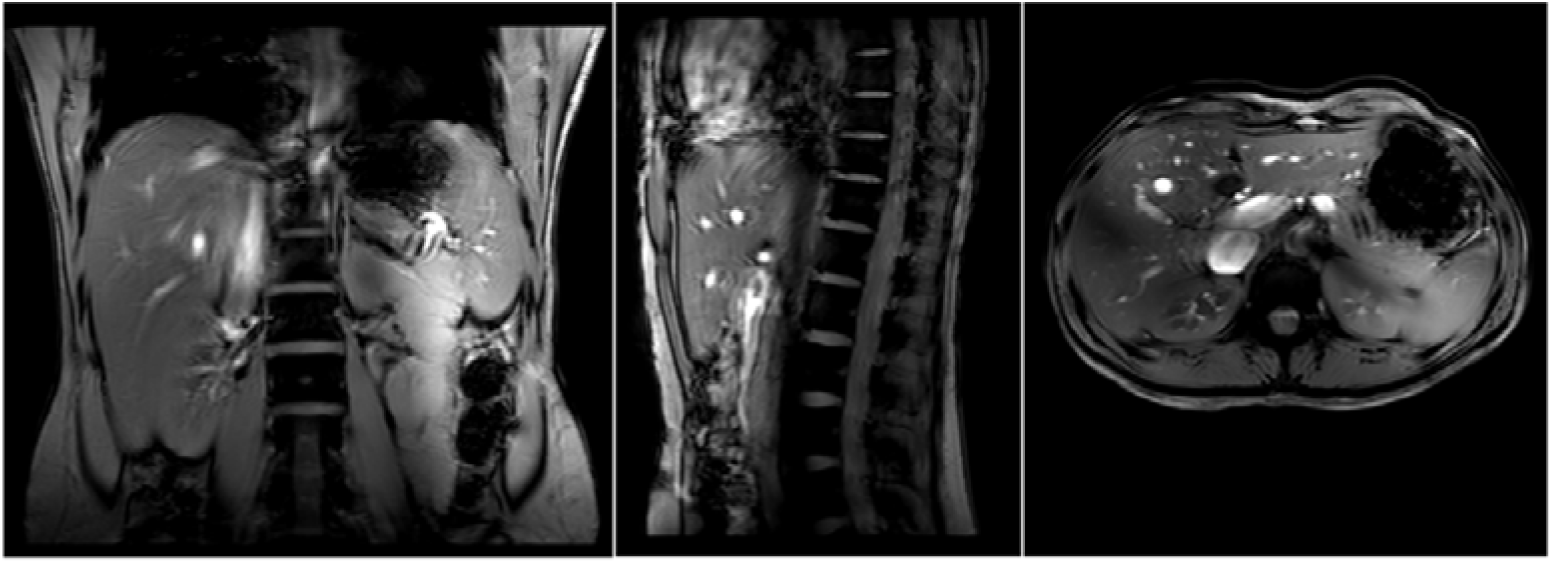
Survey of participant using eight-antennas and parallel transmit (2D gradient echo, TR 10 ms, 15o flip angle, 280 mm × 280 mm × 464 mm field of view; 257 phase encoding steps, 516 frequency encoding steps; 30 × 10-mm-thick slices, 78 s acquisition), demonstrating field of view provided by the eight antennas.

Fig. 7 shows results from a patient with liver metastases of oesophago-gastric cancer. Spectra from the liver and from the tumor allow differentiation of PE and PC. The 31P 3D CSI is projected onto a Dixon image. PE and PC appear with equal peak heights in the liver volume, where as in the tumor, the PE peak is greater than the PC peak.

**Fig. 7.**
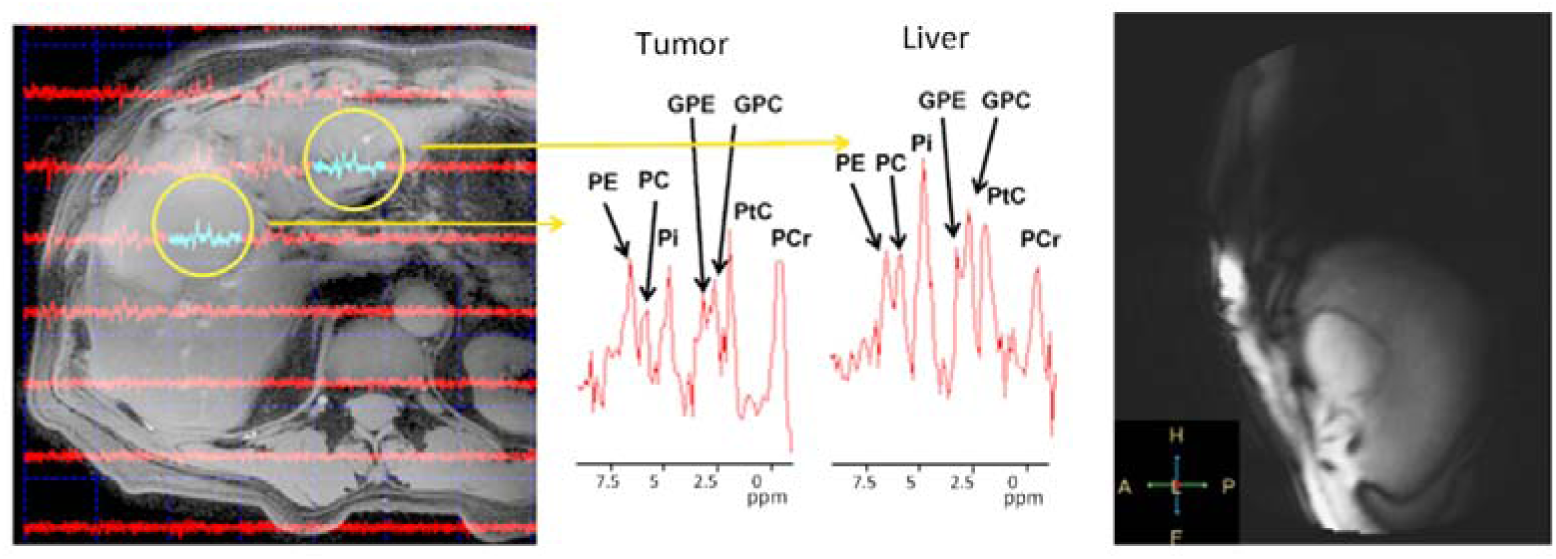
Phosphorus spectra 3DCSI from a patient with liver metastasis (gastric primary), displayed on a B1-shimmed Dixon image. PE:phosphoethanolamine, PC:phosphocholine, GPC:glycerophosphocholine, GPE: glycerophosphoethanolamine, Pi: inorganic phosphate, PtC: phosphotidylcholine, and PCr: phosphocreatine. PE of tumor is greater than PC, GPE, GPC, and Pi, while the peak height of PE is less than or equal to those of PC, GPE, GPC, and Pi in the liver voxel. The tumor is readily visible in the localizer image obtained with a pair of antennas (right).

## Discussion

We have developed a protocol for obtaining multi-nuclear and parallel transmit data and demonstrated results in a patient. The field of view of two antennas on top of the 31P loops allowed for gradient echo imaging of the liver for region-of-interest-based B0 shimming and tumor-aligned placement of the 31P 3D CSI matrix. With such a setup it is possible to locate tumors anywhere within the liver in order to optimally position the imaging matrix of 3D CSI, using the two-channel proton transmit channels available during multi-nuclear scan sessions. In the present study, the two-antenna set up failed to image the complete liver boundaries in half of the participants (four of the nine healthy volunteers, and in the patient), and there were signal drop outs within the liver for all volunteers. However, with eight-antennas, it is possible to image the entire abdomen, and to avoid image voids in the liver by B1-shimming.

In the present study, a local coil was used to excite the phosphorus signal. In order to overcome transmit inhomogeneity associated with surface coils, we combine adiabatic pulses with 3D CSI for localization as it is known to be less B1-sensitive than other methods.^24^ Breathing-belt triggering^25^ would stretch a 3D CSI single average acquisition to 25 minutes (5x increase), and was therefore deemed impractical. A limitation of the present study is that the adiabatic pulse was optimized for a uniform magnitude over the bandwidth, and not for phase which resulted in the introduction of non-linear phase offsets across the spectra. Use of a BIR4 pulse, such as that used by Runge et al. for refocusing^19^ would provide a more uniform phase across the bandwidth of the adiabatic pulse. Additionally, we can improve the penetration depth of the 31P data by optimizing the adiabatic pulse for a lower threshold B1. The TR of the present study was insufficient to overcome saturation effects in the absence of a spoiler gradient due to the widely varying longitudinal relaxation (T1) of relevant metabolites.^18,26^ Spectral quality could be improved through B1 calibration per individual and volume depth, which has the added benefit of allowing for T1-saturation correction. The use of a B1-calibration reference marker (such as in ^18^) would also facilitate quantification of metabolite concentration. Further improvements in spectral SNR are possible through implementation of various imaging techniques including AMESING,^19^ DIMEPT,^27^ and proton observed phosphorus editing.^28^ Although not widely available, hardware improvements can also be used to boost the 31P SNR. Purvis et al. use a 16-channel phosphorus receive array^18^ to acquire 31P of the liver and have demonstrated the use of a 31P birdcage for transmit in combination with a 31P receive array for cardiac studies.^29^

To our knowledge, 7T 31P methodologies for the liver have focused on diffuse liver disease using loops for 1H imaging with partial coverage of the liver (<50%).^18,19^ Full coverage of the liver is not necessary for diffuse liver disease applications. Runge et al., use two quadrature-combined overlapping 15-cm-diameter coils wrapped around the liver.^17^ Signal is concentrated within a depth of 6 cm, falling off by 8 cm, and indistinguishable from noise in some locations at 8 cm and at most 12 cm from the surface – half of the right lobe and the entire left lobe of the liver are outside of the field of view (see Fig. 1 adapted from ^19^). Purvis et al. use a 10-cm-diameter loop coil to achieve partial coverage of the left and right lobes of the liver, with signal concentrated within 6 cm of the surface of the participants, and indistinguishable from noise by 10 cm (see fig 1a and 2c in ^18^). While Runge et al., opted for a dual-tuned coil, Purvis et al., remove the 1H coil and replace it with a 31P coil array after acquiring localizer images and performing B0 shimming. By comparison, using antennas in place of loops for 1H imaging, we were able to image the left and right hemi-livers (Couinaud system) using two quadrature-combined elements, while simultaneously doubling the field of view. As antennas (1H frequency) and loops (31P frequency) are inherently decoupled,^18,30^ we did not have to either remove the 1H setup after B0 shim, or include additional circuitry (which can decrease SNR) to decouple the 1H and 31P elements.

In vitro evaluations of phospholipid metabolism in primary (hepatocellular carcinomas) and secondary (adenocarcinomas and squamous cell carcinomas) malignant tumors of the liver can clearly differentiate between tumor-containing and non-tumor-containing samples from a patient.^31^ Although in vivo 31P at lower field strengths is limited to large tumors, 31P as a marker for treatment response in hepatic primary (hepatocellular carcinomas) and liver metastases (adenocarcinomas and squamous cell carcinomas) has proven useful for identifying responders and non-responders, ^32^ even in the absence of evidence of treatment response in anatomical images. ^33^ At lower field strengths, providing reliable measures that differentiate between controls and tumors is challenging due to partial volume affects ^34^ and the overlap in the peaks of the PME’s PC and PE and the phosphodiesters (PDE) GPC and GPE due to the limited spectral resolution.^31,34^

In a recent human breast cancer study, the 31P metabolite concentrations (PE, PC, GPE, and GPE), clearly differentiated tumor tissue from healthy controls in vivo at 7T.^35^ Ultra-high field in vivo 31P spectroscopy has also been shown to relate to histological markers of aggressiveness such as mitotic count,^36^ and aid in the decision tree for determining which breast cancer patients require systemic therapy (e.g. chemotherapy or radiation).^37^ In-vivo animal models at 7T have also demonstrated that the ratio of PME to β-adenosine triphosphate (NTP) with 1.5 cm × 1.5 cm × 2 cm voxels clearly differentiated between tumor-containing volumes and liver volumes with no macroscopic tumor.^38^

There is an additional and known background signal in the liver not present in breast or prostate phosphorous spectroscopy at 2.06 ppm, corresponding to phosphatidylcholine that is present in the liver and in high concentrations in the gall bladder.^39^ As both the spectra presented in Fig. 7 come from voxels near the gall bladder, there are large peaks from phosphatidylcholine (2.06 ppm). Likewise, PCr peaks are present due to contamination from neighboring muscle tissue. The spectra from the patient with liver metastases appears to exhibit a metabolic phenotype with PE > GPE while the peak height of PC is reduced as compared to PE. The same metabolic phenotype, has been identified and correlated to aggressive cancers in NMR.^40^ However, this metabolic phenotype is as of yet not validated in vivo, and there is no consensus as to whether the phenotype observed in NMR experiments is because the preclinical models exhibiting the phenotype are not representative of the tumor micro-environment in humans—for example that the high acidity causes the GPC/PC to invert although the PE > GPE (for example in breast cancer).^41^ Further investigation within the liver and other tumor sites are warranted to explore the clinical relevance of such a metabolic phenotype.

As an improvement over conventionally used surface loops, we have demonstrated that with two-antennas combined in quadrature, we can achieve nearly full coverage of the liver, facilitating tumor localization and region-of-interest B0 shimming. While two antennas can provide near complete coverage of the liver for gradient echo scans, parallel transmit systems for B1-shimming are required for obtaining radiological grade images that require uniform contrast weighting in T1 and T2 weighted MRI. Through the use of antennas for proton imaging and parallel transmit in combination with 31P coil loops, we can begin to explore the potential of adapting treatments to patients using full-radiological grade images and multi-parametric imaging.

## References

1. Glunde K, Jiang L, Moestue SA, Gribbestad IS. MRS and MRSI guidance in molecular medicine: targeting and monitoring of choline and glucose metabolism in cancer. NMR Biomed. 2011;24(6):673-690.

2. Disibio G, French SW. Metastatic patterns of cancers: results from a large autopsy study. Arch Pathol Lab Med. 2008;132(6):931-939.

3. Pentheroudakis G, Fountzilas G, Bafaloukos D, et al. Metastatic breast cancer with liver metastases: a registry analysis of clinicopathologic, management and outcome characteristics of 500 women. Breast Cancer Res Treat. 2006;97(3):237-244.

4. Wagner AD, Unverzagt S, Grothe W, et al. Chemotherapy for advanced gastric cancer. Cochrane Database Syst Rev. 2010(3):CD004064.

5. Golubnitschaja O, Sridhar KC. Liver metastatic disease: new concepts and biomarker panels to improve individual outcomes. Clin Exp Metastasis. 2016;33(8):743-755.

6. Cheng M, Bhujwalla ZM, Glunde K. Targeting Phospholipid Metabolism in Cancer. Front Oncol. 2016;6:266.

7. Podo F, Canevari S, Canese R, Pisanu ME, Ricci A, Iorio E. MR evaluation of response to targeted treatment in cancer cells. NMR Biomed. 2011;24(6):648-672.

8. Klomp DW, van de Bank BL, Raaijmakers A, et al. 31P MRSI and 1H MRS at 7 T: initial results in human breast cancer. NMR Biomed. 2011;24(10):1337-1342.

9. Dietz C, Ehret F, Palmas F, et al. Applications of high-resolution magic angle spinning MRS in biomedical studies II-Human diseases. NMR Biomed. 2017;30(11).

10. Metzger GJ, Snyder C, Akgun C, Vaughan T, Ugurbil K, Van de Moortele PF. Local B1+ shimming for prostate imaging with transceiver arrays at 7T based on subject-dependent transmit phase measurements. Magn Reson Med. 2008;59(2):396-409.

11. Vaughan JT, Snyder CJ, DelaBarre LJ, et al. Whole-body imaging at 7T: preliminary results. Magn Reson Med. 2009;61(1):244-248.

12. Pang Y, Wu B, Wang C, Vigneron DB, Zhang X. Numerical Analysis of Human Sample Effect on RF Penetration and Liver MR Imaging at Ultrahigh Field. Concepts Magn Reson Part B Magn Reson Eng. 2011;39B(4):206-216.

13. Roschmann P. Radiofrequency penetration and absorption in the human body: limitations to high-field whole-body nuclear magnetic resonance imaging. Med Phys. 1987;14(6):922-931.

14. Raaijmakers AJ, Ipek O, Klomp DW, et al. Design of a radiative surface coil array element at 7 T: the single-side adapted dipole antenna. Magn Reson Med. 2011;66(5):1488-1497.

15. Raaijmakers AJ, Luijten PR, van den Berg CA. Dipole antennas for ultrahigh-field body imaging: a comparison with loop coils. NMR Biomed. 2016;29(9):1122-1130.

16. Van de Moortele PF, Akgun C, Adriany G, et al. B(1) destructive interferences and spatial phase patterns at 7 T with a head transceiver array coil. Magn Reson Med. 2005;54(6):1503-1518.

17. Padormo F, Beqiri A, Hajnal JV, Malik SJ. Parallel transmission for ultrahigh-field imaging. NMR Biomed. 2016;29(9):1145-1161.

18. Purvis LAB, Clarke WT, Valkovic L, et al. Phosphodiester content measured in human liver by in vivo (31) P MR spectroscopy at 7 tesla. Magn Reson Med. 2017;78(6):2095-2105.

19. Runge JH, van der Kemp WJ, Klomp DW, Luijten PR, Nederveen AJ, Stoker J. 2D AMESING multi-echo (31)P-MRSI of the liver at 7T allows transverse relaxation assessment and T2-weighted averaging for improved SNR. Magn Reson Imaging. 2016;34(2):219-226.

20. Raaijmakers AJ, Italiaander M, Voogt IJ, et al. The fractionated dipole antenna: A new antenna for body imaging at 7 Tesla. Magn Reson Med. 2016;75(3):1366-1374.

21. Hoogduin H RA, Visser F, Luijten P (2014) In: Proceedings of the 22nd scientific meeting Initial experience with BOLD imaging of the kidneys at 7 T. International Society for Magnetic Resonance in Medicine; 2014; Milan.

22. Dixon WT. Simple proton spectroscopic imaging. Radiology. 1984;153(1):189-194.

23. Naressi A, Couturier C, Devos JM, et al. Java-based graphical user interface for the MRUI quantitation package. MAGMA. 2001;12(2-3):141-152.

24. Gonen O, Murphy-Boesch J, Li CW, Padavic-Shaller K, Negendank WG, Brown TR. Simultaneous 3D NMR spectroscopy of proton-decoupled fluorine and phosphorus in human liver during 5-fluorouracil chemotherapy. Magn Reson Med. 1997;37(2):164-169.

25. Barbieri S, Donati OF, Froehlich JM, Thoeny HC. Comparison of Intravoxel Incoherent Motion Parameters across MR Imagers and Field Strengths: Evaluation in Upper Abdominal Organs. Radiology. 2016;279(3):784-794.

26. Chmelik M, Povazan M, Krssak M, et al. In vivo (31)P magnetic resonance spectroscopy of the human liver at 7 T: an initial experience. NMR Biomed. 2014;27(4):478-485.

27. van der Kemp WJ, Boer VO, Luijten PR, Klomp DW. Increased sensitivity of 31P MRSI using direct detection integrated with multi-echo polarization transfer (DIMEPT). NMR Biomed. 2014;27(10):1248-1255.

28. Wijnen JP, Klomp DW, Nabuurs CI, et al. Proton observed phosphorus editing (POPE) for in vivo detection of phospholipid metabolites. NMR Biomed. 2016;29(9):1222-1230.

29. Valkovic L, Dragonu I, Almujayyaz S, et al. Using a whole-body 31P birdcage transmit coil and 16-element receive array for human cardiac metabolic imaging at 7T. PLoS One. 2017;12(10):e0187153.

30. Rivera DS, Wijnen JP, van der Kemp WJ, Raaijmakers AJ, Luijten PR, Klomp DW. MRI and (31)P magnetic resonance spectroscopy hardware for axillary lymph node investigation at 7T. Magn Reson Med. 2015;73(5):2038-2046.

31. Cox IJ, Bell JD, Peden CJ, et al. In vivo and in vitro 31P magnetic resonance spectroscopy of focal hepatic malignancies. NMR Biomed. 1992;5(3):114-120.

32. Dixon RM, Angus PW, Rajagopalan B, Radda GK. Abnormal phosphomonoester signals in 31P MR spectra from patients with hepatic lymphoma. A possible marker of liver infiltration and response to chemotherapy. Br J Cancer. 1991;63(6):953-958.

33. Meyerhoff DJ, Karczmar GS, Valone F, Venook A, Matson GB, Weiner MW. Hepatic cancers and their response to chemoembolization therapy. Quantitative image-guided 31P magnetic resonance spectroscopy. Invest Radiol. 1992;27(6):456-464.

34. Francis IR, Chenevert TL, Gubin B, et al. Malignant hepatic tumors: P-31 MR spectroscopy with one-dimensional chemical shift imaging. Radiology. 1991;180(2):341-344.

35. Wijnen JP, van der Kemp WJ, Luttje MP, Korteweg MA, Luijten PR, Klomp DW. Quantitative 31P magnetic resonance spectroscopy of the human breast at 7 T. Magn Reson Med. 2012;68(2):339-348.

36. Schmitz AM, Veldhuis WB, Menke-Pluijmers MB, et al. Multiparametric MRI With Dynamic Contrast Enhancement, Diffusion-Weighted Imaging, and 31-Phosphorus Spectroscopy at 7 T for Characterization of Breast Cancer. Invest Radiol. 2015;50(11):766-771.

37. Schmitz AMT, Veldhuis WB, Menke-Pluijmers MBE, et al. Preoperative indication for systemic therapy extended to patients with early-stage breast cancer using multiparametric 7-tesla breast MRI. PLoS One. 2017;12(9):e0183855.

38. McKenzie EJ, Jackson M, Sun J, Volotovskyy V, Gruwel ML. Monitoring the development of hepatocellular carcinoma in woodchucks using 31P-MRS. MAGMA. 2005;18(4):201-205.

39. Chmelik M, Valkovic L, Wolf P, et al. Phosphatidylcholine contributes to in vivo (31)P MRS signal from the human liver. Eur Radiol. 2015;25(7):2059-2066.

40. Moestue SA, Borgan E, Huuse EM, et al. Distinct choline metabolic profiles are associated with differences in gene expression for basal-like and luminal-like breast cancer xenograft models. BMC Cancer. 2010;10:e433.

41. Haukaas TH, Euceda LR, Giskeodegard GF, Bathen TF . Metabolic Portraits of Breast Cancer by HR MAS MR Spectroscopy of Intact Tissue Samples. Metabolites. 2017;7(2).

